# Introduce a New Approach to Detect Genes Associated to Oral Squamous Cell Carcinoma

**DOI:** 10.1101/377788

**Authors:** Jianqiang Li, Caiyun Yang, Yang Ji-Jiang, Shi Chen, Qing Wang, HuiPan, Siyuan Liang, Weiliang Qiu

**Affiliations:** Faculty of Information Technology, Beijing University of Technology, Beijing, 100124, China; Beijing Engineering Research Center for IoT Software and Systems, Beijing, 100124, China; Tsinghua National Laboratory for Information Science and Technology, Tsinghua University, Beijing, 100084, China; Department of Endocrinology, Peking Union Medical College Hospital, Chinese Academy of Medical Sciences & Peking Union Medical College, Beijing, 100730, China; Channing Division of Network Medicine, Brigham and Women’s Hospital/Harvard Medical School, 181 Longwood Avenue, Boston MA 02115, USA

**Keywords:** missing heritability, differential variability, KEGG

## Abstract

Oral squamous cell carcinoma (OSCC) represents the most frequent of all oral neoplasms in the world. Genetics plays an important role in the etiopathogenesis of OSCC. However, the investigation of the molecular mechanism of OSCC is still incomplete. In this article, we introduced a new approach to detect OSCC-associated genes, in which we not only compare mean difference, but also variance difference between cases and controls. Based on two OSCC datasets from Gene Expression Omnibus, we identified 456 differentially variable (DV) gene probes, in addition to 2,375 differentially expressed (DE) gene probes. There are 2,193 DE-only probes, 274 DV-only probes, and 182 DE-and-DV probes. DAVID functional analysis showed that genes corresponding to DE-only, DV-only, and DE-and-DV probes were enriched in different KEGG pathways, indicating they play different roles in OSCC. This new approach can be used to investigate the genetic risk factors for other complex human diseases.

## Introduction

Oral squamous cell carcinoma (OSCC), the most frequent of all oral neoplasms in the world, is considerably significant for public health [1, 2]. Studies show that OSCC is associated with a mass of mortality and morbidity [3]. OSCC is a multistep progression, which is influenced by several environmental factors such as tobacco chewing, smoking, alcohol consumption, and infection of human papilloma virus (HPV) [4]. Genetic alterations, such as the change of gene expressions of tumor suppressor genes and oncogenes, also play important roles in the development of OSCC [5]. A few review articles [6-9] summarized genetic anomalies associated with OSCC according to the type of structure affected (chromosome, allele, oncogene, tumor suppressor gene, or nucleotide) and the type of anomaly (polymorphism, point mutation, deletion, and other alterations). Many OSCC-associated genes (e.g., TP53, CDKN2A, Cyclin D1, EGFR, SMAD-2, SMAD-4, CDK2AP1, FHIT) have been identified [4, 7, 8, 10].

However, the investigation of the molecular mechanism of OSCC is still incomplete. The efforts to identify novel OSCC-associated genes are continuing [11-13].

In fact, the “missing heritability” problem has not been solved yet since Manolio et al. (2009) [14] raised the problem that genetic variants in Genome-Wide Association Study (GWAS) cannot completely explain the heritability of complex traits [15]. Several mechanisms have been proposed to account for the gap in heritability, such as epigenetics, epistasis, sequencing depth, and microbiome [15]. Recent years, researchers have also investigated non-coding genome and splice sites [16]. New methods [17, 18] have also been proposed to obtain a more accurate estimate of heritability. In this article, we hypothesized that (1) more genes are associated with OSCC and (2) identifying more OSCC-associated genes would help reduce the gap in OSCC heritability. To identify novel genes associated with OSCC, we introduced in this article a new method based on the observation that existing methods to identify OSCC-associated genes focus on comparing the mean expression levels of genes between OSCC samples and control samples. Variance is another important measurement in Statistics and indicates how much information a dataset contains. The larger the variance is, the more information the dataset contains.

Several studies [19–21] have demonstrated that differential variable epigenetic marks (specifically, DNA methylation marks) are risk factors for complex human diseases. For gene expression data, there are also several papers [22–26] investigated differential variable gene probes. Ho et al. (2008) demonstrated that genes differentially variable (i.e., having different variances of expression levels) between two biological states (i.e., disease vs. non-disease) are also biologically relevant [22].

Strbenac et al. (2016) proposed to detect DE, DV, and differential distributed (DD) gene probes. Their methods can work for both microarray gene expression data and RNA-seq data. However, Strbenac et al. (2016) did not check the enriched pathways for DE-only, DV-only, and DE-and-DV genes. Ran and Daye (2017) proposed a generalized linear model to fit RNA-seq count data to detect DE genes and DV genes. They reported enriched pathways for DE-only genes and for DV-only genes, but not for DE-and-DV genes. Rahmatallah et al. (2017) and Sanati et al. (2018) considered DE, DV, and differentially correlated (DC) genes, separately. Their methods can work for both microarray gene expression data and RNA-seq data. However, Rahmatallah et al. (2017) and Sanati et al. (2018) focused on DC genes and did not try to distinguish DE-only, DV-only, and DE-and-DV genes. All the aforementioned 5 papers have not been applied to detect DV genes for OSCC. In this article, we proposed to identify 3 sets of OSCC-associated genes: (1) genes corresponding to gene probes that are only differentially expressed between OSCC samples and non-OSCC samples; (2) genes corresponding to gene probes that are only differentially variable (i.e., having different variances) between OSCC samples and non-OSCC samples; and (3) genes corresponding to gene probes that are both differentially expressed and differentially variable between OSCC samples and non-OSCC samples.

## Materials and Methods

### Two Datasets

In this paper, we aimed to detect genes associated with OSCC based on two gene microarray datasets that are available in the Gene Expression Omnibus (GEO) [27]: GSE30784 (the discovery set) (https://www.ncbi.nlm.nih.gov/gquery/?term=GSE30784) and GSE6791 (the validation set) (https://www.ncbi.nlm.nih.gov/gquery/?Term=GSE6791). The GSE30784 dataset contains 167 OSCC cancer cases, 17 dysplasia samples, and 45 controls [28]. In our study, we excluded the 17 dysplasia samples and only compared the 167 OSCC cancer cases with the 45 controls. The GSE6791 dataset includes 42 head and neck squamous cell carcinoma (HNSCC) cases, 14 head and neck normals (HNN), 20 cervical cancers, and 8 cervical normal [29]. In our study, we excluded the 28 cervical samples and only compared the 42 HNSCC cases and the 14 controls (HNN). We used GSE30784 as the discovery set and GSE6791 as the validation set. We chose GSE6791 as the validation set because 1) both GSE30784 and GSE6791 were generated using the same Affymetrix U133 Plus 2.0 GeneChip platform; 2) HNSCC arises from the mucosal surfaces of the oral cavity (OSCC), oropharynx (OPSCC) and larynx; and 3) OSCC shares some of the same risk factors as HNSCC, such as cigar and pipe smoking [30]. For each sample, the expression levels of 54,675 probes were measured by using Affymetrix Human Genome U133 Plus 2.0 Array.

### Data Quality Check

It is a routine to check the data quality before the statistical analysis of gene expression data. The expression data in GSE30784 and GSE6791 have been log2 transformed and quantile normalized (i.e., they are not raw expression data). We double-checked data quality to identify any gene probes with outlying expression, any arrays with low quality, and any batch effects.

For GSE30784 and GSE6791, we kept the same set of 39,534 consensus probes, which are nucleotide sequences assembled by Affymetrix, based on one or more sequence taken from a public database (http://www.ebi.ac.uk/arrayexpress/arrays/A-GEOD-13158/?ref=E-MTAB-2024), hence are more likely from the genome. We excluded 17 dysplasia samples from GSE30784 and 28 cervical samples from GSE6791. To check if there are batch effects or outlying probes or arrays, we drew the plots of quantiles of expression levels across arrays (Suppl.Fig.1) and the PCA plots (plots of the first principal component versus the second principal components) (Suppl.Fig.2) for both datasets. No obvious outlying probes or arrays were found in these plots. No obvious technical batch effects were found, except for the effect of cancer status.

Next, we eliminated the probes that either showed no variation across the samples (inter quartile range (IQR) of expression levels less than 0.1 on log2 scale) or were expressed at very low magnitude (any probe in which the maximum expression value for that probe in any of the samples was less than 3 on log2 scale) [28]. 20,683 probes were kept in GSE30784 and 39,506 probes were kept in GSE6791. We also excluded the probes without annotations of gene symbols or Entrez ids for two gene datasets. Finally, we used 10,754 consensus probes, which were appeared in both the discovery set and the validation set, for the downstream analyses.

### Differential Variability Analysis

Differential variability (DV) analysis aims to identify genes with a significant change in the variance of expression between diseased patients and non-diseased individuals (e.g., cancer cases and controls) [22]. For a given probe, denote 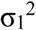 and 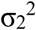 as the variances of gene expression for cases and controls, respectively. We would like to test the null hypothesis that the variances are equal 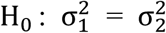 versus the alternative hypothesis that the variances are different 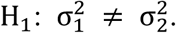

F test, comparing the sample variances of the two groups, is the most commonly used statistical test for the equal variance [31]. However, F test is very sensitive to the violation of the normality assumption (i.e., assuming data from cases and controls both follow normal distributions). Many tests robust to the violation of the normality assumption have been proposed. The Brown-Forsythe test (BF test) is one of the robust equal-variance tests, which results from an ordinary one-way analysis of variance based on the absolute deviations from the group median [32]. The BF test appears to be one of the best equal-variance tests in terms of controlling type I error rate and having large testing power, based on the systematic simulation studies concluded by Conover et al. [33]. In this article, we used the BF test to identify gene probes having significantly different variances of gene expression between two conditions：OSCC cancer versus control.

### Statistical Analysis

We used the 10,754 gene probes and 212 samples (167 OSCC cancer samples and 45 control samples) in GSE30784 as the discovery set, and used the same 10,754 probes and 54 samples (42 HNSCC samples and 14 HNN samples) in GSE6791 as the validation set. To identify differentially expressed (DE) genes, we performed moderated t-test adjusting for age and sex by using R Bioconductor package *limma* [34]. To identify differentially variable (DV) genes, for each of the 10,754 probes in the discovery set (GSE30784), we performed the BF test. To control for multiple testing, we used Benjamini and Hochberg’s method to adjust for p-values of DE tests (DV tests) so that the false discovery rate (FDR) < 0.05 [35]. We claimed a probe in the discovery set (GSE30784) as a DE probe (DV probe) if its FDR-adjusted p-value < 0.05 in DE tests (DV tests). For the detected DE (DV) probes in the discovery set (GSE30784), we performed the moderated t-test (BF test) based on the validation set (GSE6791). If the un-adjusted p-value for the probe is < 0.05 in the validation set (GSE6791) and if the sign of m_1_-m_2_ 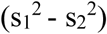 is the same in the discovery set as in the validation set, we claimed that the DE (DV) probe detected in the discovery set (GSE30784) is validated in the validation set (GSE6791), where m_1_ and 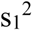 are the sample mean and sample variance for the cases and m_2_ and 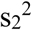 are the sample mean and variance for the controls.

### Pathway Enrichment Analysis

To distinguish the contributions of the validated DE probes and the validated DV probes on the mechanisms of OSCC, we first classified the validated probes to 3 groups: (1) DE-only probes (validated DE probes, but not validated DV probes); (2) DV-only probes (validated DV probes, but not validated DE probes); and (3) DE-and-DV probes (validated DE probes and validated DV probes). We then obtained the genes corresponding to each of the 3 groups of validated probes. We next did the KEGG pathway enrichment analysis by DAVID 6.8 [36] for each of the three groups of genes.

## Results

### The DE gene probes

Through DE analysis we got 6,239 significant DE gene probes (FDR adjusted p-value <0.05) based on the discovery set GSE30784, 2,375 of which were validated in the validation set (GSE6791). Suppl.Table1 lists the 2,375 validated DE probes. Fig.1 shows the parallel boxplots of gene expression levels for the top 2 validated DE probes. Denote the number of the validated DE gene probes as n.validated and the number of DE probes in the discovery set as n.dis. The proportion n.validated/n.dis is 38.07%.

**Fig.1:**
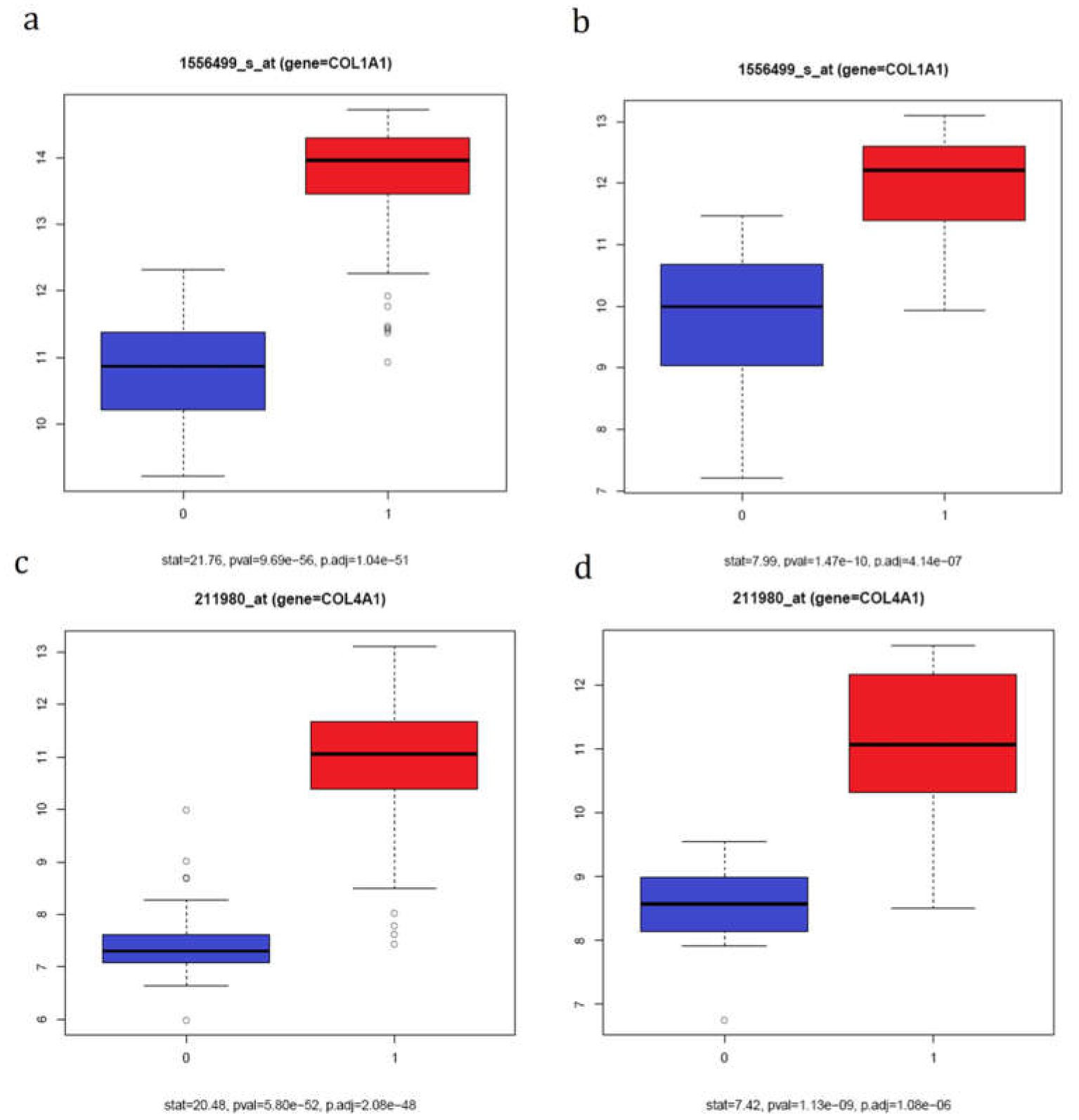
Parallel boxplots of log2 gene expression between cancer samples (1) and control samples (0) for the most and second most significant validated DE gene probes. The x-axis indicates if a sample is a cancer sample (1) or control sample (0). The y-axis is the log2 gene expression levels. Subfigures a and c are based on the discovery set GSE30784. Subfigure b and d are based on the validation set GSE6791.

### The DV gene probes

For the discovery set GSE30784, we performed the BF test for each of the 10,754 probes, and got 2,904 significant DV gene probes (FDR adjusted p-value <0.05), among which 456 probes were validated in the validation set (GSE6791), Suppl.Table2 lists the 456 validated DV probes. Fig.2 shows the parallel boxplots of gene expression levels for the top 2 validated DV probes. Denote the number of the validated DV gene probes as n.validated and the number of DV probes in the discovery set as n.dis. The proportion n.validated/n.dis is 15.70%, which is significantly larger than what we expect by chance.

**Fig.2:**
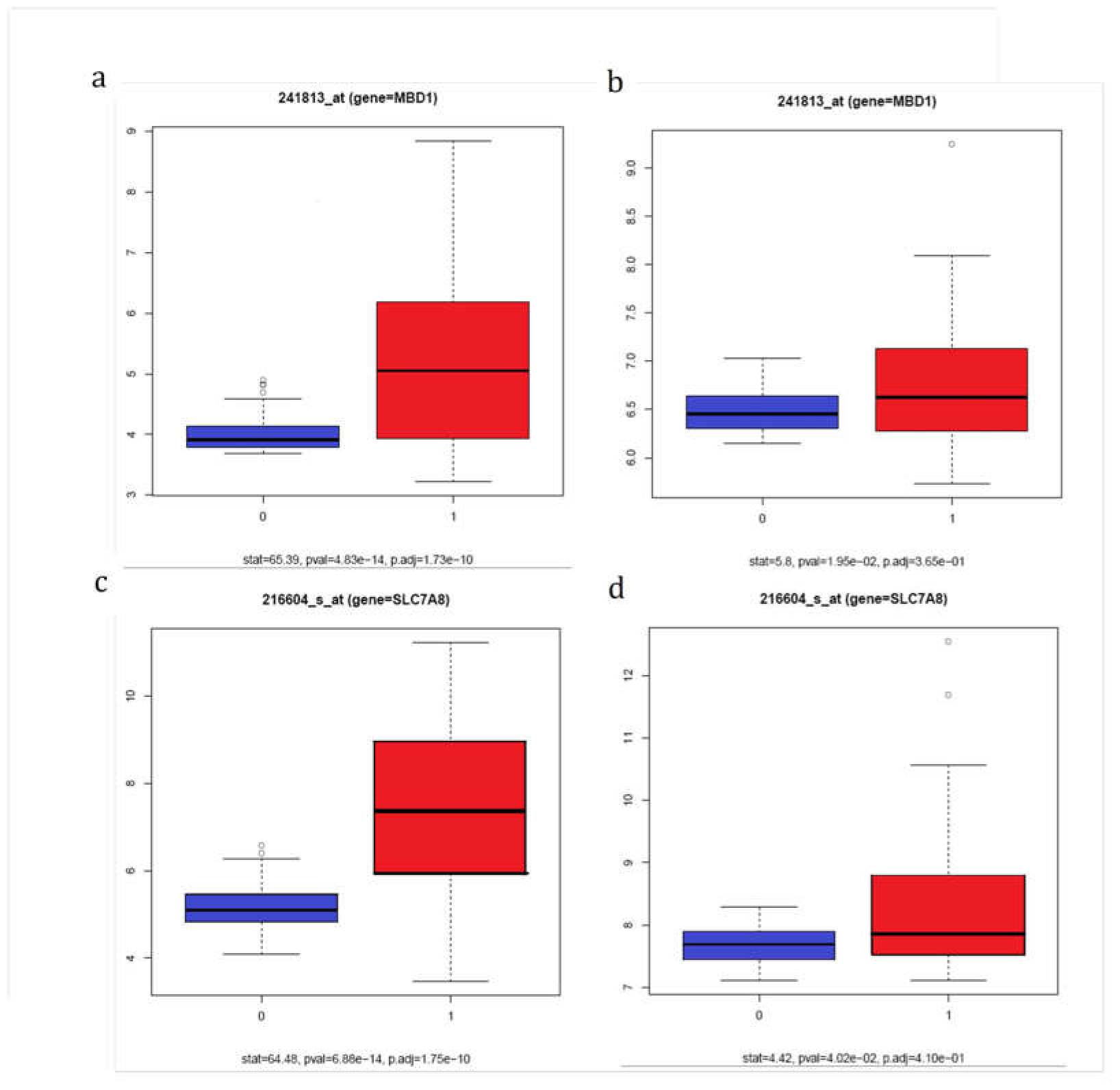
Parallel boxplots of log2 gene expression between cancer samples (1) and control samples (0) for the most and second most significant validated DV gene probes. The x-axis indicates if a sample is a cancer sample (1) or control sample (0). The y-axis is the log2 gene expression levels. Subfigures a and c are based on the discovery set GSE30784. Subfigure b and d are based on the validation set GSE6791.

### DE-only gene probes, DV-only gene probes and DE-and-DV gene probes

Among the 2,375 validated DE gene probes, 2,193 probes were only differentially expressed, not differentially variable (Suppl.Table.3). Table1 lists the top 10 validated DE-only gene probes. Among the 456 validated DV gene probes, 274 probes were only differentially variable, not differentially expressed (Suppl.Table.4). Table2 lists the top 10 validated DV only gene probes. Fig.3 shows the Venn diagram of the 2 groups of probes (the validated DE probes and the validated DV probes). Suppl. Fig.3 shows the parallel boxplots of gene expression levels for the top 2 DE-and-DV probes.

**Table1:** Information about the top 10 validated DE-only gene probes. This table is in ascending order of pval.DE (i.e., the p-value to test for equal mean based on the discovery set).

| probe | gene | stat.DE | pval.DE | p.adj.DE | stat.valid | pval.valid |
| --- | --- | --- | --- | --- | --- | --- |
| 1556499_s_at | COL1A1 | 21.76 | 9.69E-56 | 1.04E-51 | 7.99 | 1.47E-10 |
| 211980_at | COL4A1 | 20.48 | 5.80E-52 | 2.08E-48 | 7.42 | 1.13E-09 |
| 241233_x_at | ANKRD20A11P | -20.48 | 5.81E-52 | 2.08E-48 | -7.62 | 5.64E-10 |
| 211964_at | COL4A2 | 18.08 | 1.07E-44 | 2.30E-41 | 7.08 | 3.93E-09 |
| 215076_s_at | COL3A1 | 17.49 | 7.13E-43 | 1.09E-39 | 6.45 | 3.89E-08 |
| 230740_at | EHD3 | -17.4 | 1.33E-42 | 1.78E-39 | -4.54 | 3.41E-05 |
| 227969_at | PCBP1-AS1 | -17.34 | 2.06E-42 | 2.46E-39 | -3.78 | 4.10E-04 |
| 212236_x_at | JUP | 16.84 | 7.55E-41 | 8.12E-38 | 6.85 | 9.05E-09 |
| 212012_at | PXDN | 16.39 | 1.98E-39 | 1.78E-36 | 5.05 | 5.94E-06 |
| 224954_at | MIR6778 | -16.3 | 3.62E-39 | 2.99E-36 | -6.27 | 7.45E-08 |

**Table2:** Information about the top 10 validated DV-only gene probes. This table is in ascending order of pval.DV (i.e., the p-value to test for equal variance based on the discovery set).

| probe | gene | stat.DV | pval.DV | p.adj.DV | stat.valid | pval.valid |
| --- | --- | --- | --- | --- | --- | --- |
| 241813_at | MBD1 | 65.39 | 4.83E-14 | 1.73E-10 | 5.8 | 1.95E-02 |
| 216604_s_at | SLC7A8 | 64.48 | 6.88E-14 | 1.75E-10 | 4.42 | 4.02E-02 |
| 52255_s_at | COL5A3 | 49.84 | 2.40E-11 | 2.87E-08 | 4.82 | 3.24E-02 |
| 1554795_a_at | FBLIM1 | 41.83 | 6.87E-10 | 4.62E-07 | 8.23 | 5.88E-03 |
| 230381_at | C1orf186 | 37.46 | 4.52E-09 | 1.80E-06 | 7.45 | 8.55E-03 |
| 238673_at | SAMD12 | 37.34 | 4.76E-09 | 1.83E-06 | 7.26 | 9.40E-03 |
| 230722_at | BNC2 | 36.49 | 6.90E-09 | 2.32E-06 | 4.85 | 3.19E-02 |
| 235683_at | SESN3 | 35.69 | 9.77E-09 | 3.00E-06 | 12.78 | 7.49E-04 |
| 226930_at | FNDC1 | 32.28 | 4.41E-08 | 1.01E-05 | 5.67 | 2.08E-02 |
| 221900_at | COL8A2 | 32.23 | 4.51E-08 | 1.01E-05 | 12.61 | 8.04E-04 |

**Fig.3:**
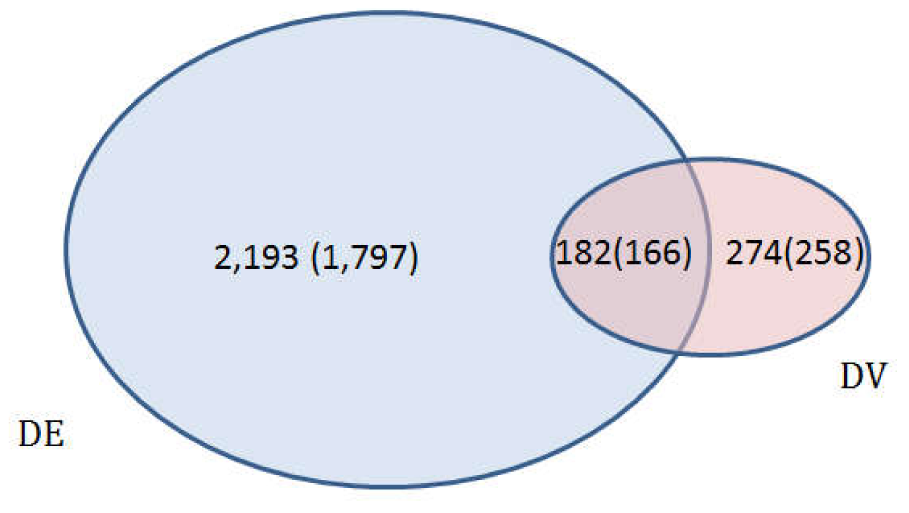
Venn diagram of the validated DE probes and validated DV probes. The left blue oval represents the set of the 2,193 DE-only gene probes (corresponding to 1,797 genes). The right red oval represents the set of the 274 DV-only gene probes (corresponding to 258 genes). There are 182 DE-and-DV probes (corresponding to 166 genes).

### DAVID enrichment analysis and GeneMANIA analysis

We performed KEGG pathway analysis for the 1,797 genes corresponding to the 2,193 validated DE-only gene probes, the 166 genes corresponding to the 182 DE-and-DV gene probes and the 258 genes corresponding to the 274 DV-only probes. For the 1,797 genes corresponding to the 2,193 validated DE only genes, there are ten significantly enriched pathways (FDR < 0.05): Focal adhesion, Protein processing in endoplasmic reticulum, Adherens junction, Epstein-Barr virus infection, Chronic myeloid leukemia, Renal cell carcinoma, Proteoglycans in cancer, Ubiquitin mediated proteolysis, Bacterial invasion of epithelial cells and Hepatitis C (Table3). For the 166 genes corresponding to the 182 DE-and-DV probes, there are two significantly enriched pathways (FDR < 0.05): ECM-receptor interaction and Amoebiasis (Table4). There are no overlapping enriched pathways. For the 258 genes corresponding to the 274 validated DV-only probes, DAVID did not report any significantly enriched pathways. However, GeneMANIA [37] analysis showed that the 258 genes are densely connected based on knowledge reported in the literature, such as co-expressed and physically interacted (Fig.4).

**Table3:** Information about the significantly enriched pathways for the 1,797 genes corresponding to the 2,193 validated DE only gene probes. This table is in ascending order of the p-value.

| Term | Count | P-Value | Benjamini |
| --- | --- | --- | --- |
| Focal adhesion | 42 | 4.80E-07 | 1.30E-04 |
| Protein processing in endoplasmic reticulum | 36 | 1.30E-06 | 1.70E-04 |
| Adherens junction | 19 | 3.10E-05 | 2.80E-03 |
| Epstein-Barr virus infection | 34 | 1.10E-04 | 7.80E-03 |
| Chronic myeloid leukemia | 18 | 1.30E-04 | 7.20E-03 |
| Renal cell carcinoma | 16 | 4.30E-04 | 1.90E-02 |
| Proteoglycans in cancer | 33 | 6.70E-04 | 2.50E-02 |
| Ubiquitin mediated proteolysis | 25 | 8.50E-04 | 2.80E-02 |
| Bacterial invasion of epithelial cells | 17 | 1.10E-03 | 3.30E-02 |
| Hepatitis C | 24 | 1.30E-03 | 3.40E-02 |

**Table4:** Information about the significantly enriched pathway for the 166 genes corresponding to gene probes that are both validated DE gene probes and validated DV gene probes. This table is in ascending order of the p-value.

| Term | Count | P-Value | Benjamini |
| --- | --- | --- | --- |
| ECM-receptor interaction | 8 | 7.00E-06 | 8.40E-04 |
| Amoebiasis | 7 | 2.40E-04 | 1.40E-02 |

**Fig.4:**
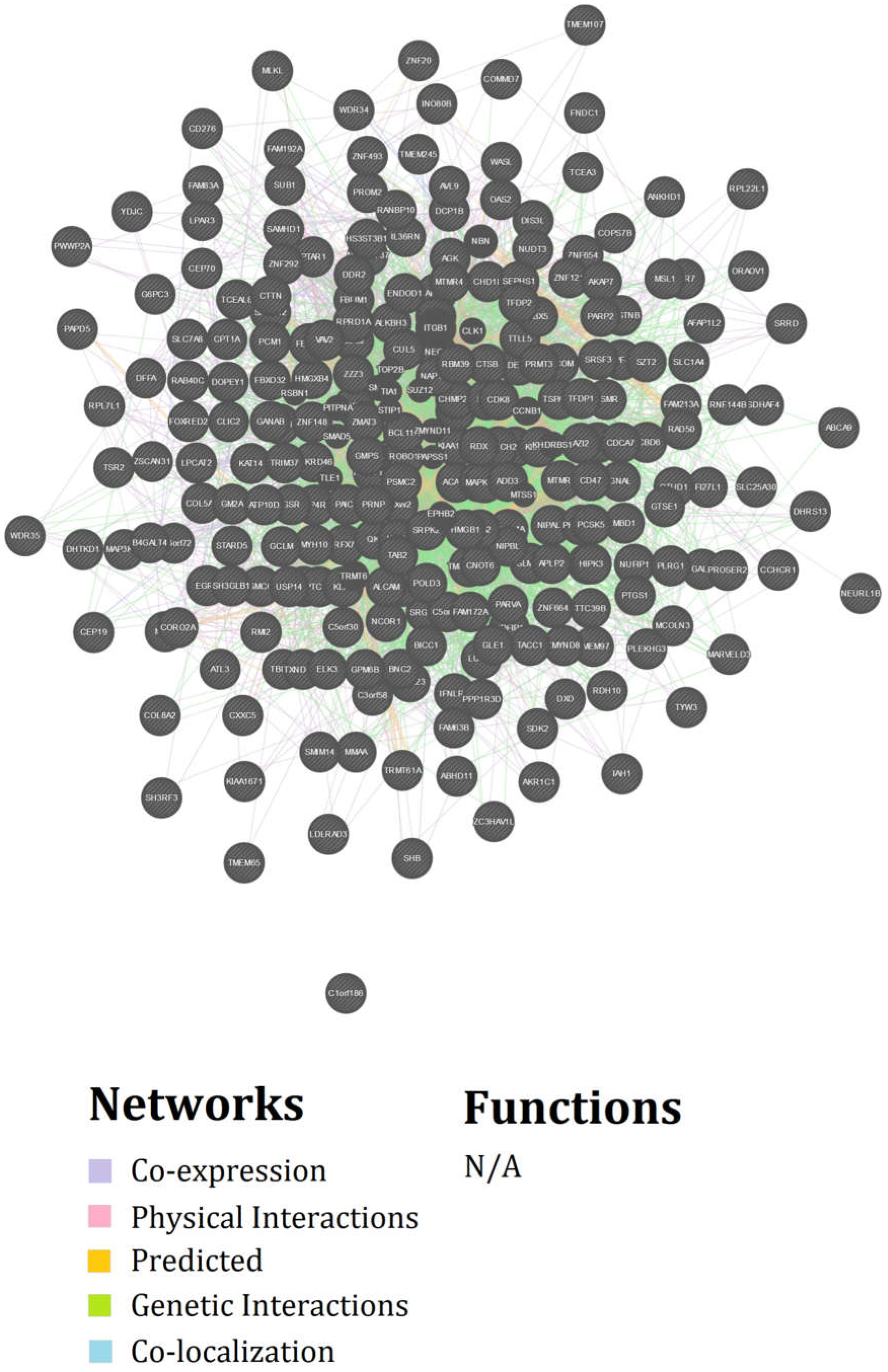
Networks of the 258 genes corresponding to the 274 DV-only gene probes obtained by GeneMania.

## Discussion

In this paper, we aimed to detect novel OSCC-associated genes by considering both mean difference and variance difference of gene expression between OSCC patients and healthy controls based on two gene microarray datasets. We obtained 3 sets of genes (1,797 genes corresponding to 2,193 DE-only probes, 258 genes corresponding to 274 DV-only probes, and 166 genes corresponding to 182 DE-and-DV probes) and found these 3 sets of genes contribute differently to OSCC in terms of KEGG pathways where they enriched.

DAVID pathway enrichment analysis showed that the 166 genes corresponding to the 182 DE-and-DV gene probes were enriched in the ECM-receptor interaction pathway, and amoebiasis. Osathanon et al. (2016) and Ge et al. (2015) both reported that the two enriched pathways are related to OSCC [38, 39]. Li et al. (2017) reported that the ECM-receptor interaction was one significant pathway related to OSCC [40]. The extracellular matrix (ECM), consisting of a complex mixture of structural and functional macromolecules plays an important role in tissue and organ morphogenesis and the maintenance of cell and tissue structure and function [41]. The 1,797 genes corresponding to the 2,193 DE-only gene probes are enriched in 10 KEGG pathways. These included focal adhesion, protein processing in endoplasmic reticulum, chronic myeloid leukemia, renal cell carcinoma, ubiquitin mediated proteolysis and hepatitis C, which have been previously reported in OSCC literature [38-40, 42]. Adherens junction was associated with the transformation from dysplasia to OSCC [43].

Epstein-Barr virus infection was reported statistically associated with increased risk of OSCC [44]. Peng et al found that copy number alterations in OSCC were distributed in the proteoglycan metabolism pathway [45]. The Bacterial invasion of epithelial cells pathway has not been reported its relation with OSCC yet.

For the 258 genes corresponding to the 274 DV-only gene probes, no significantly enriched pathways were found. However, GeneMANIA analysis showed that there are many co-expressions, physical interactions, predicted, genetic interactions and co-localization among the 258 genes based on existing literature. We also found that seven out of the ten genes corresponding to the top ten of the 274 DV only gene probes (i.e., MBD1, SLC7A8, COL5A3, FBLIM1, C1orf186, SAMD12, BNC2, SESN3, FNDC1, and COL8A2), are related to OSCC. He et al. (2015) reported that MBD1 plays an important role in cell migration and invasion in OSCC [46]. SLC7A8 gene was in a gene network in the OSCC cell lines, which is related to cancer, cellular movement, and cell [47]. Sun et al. (2014) reported that COL5A3 was in the KEGG pathway hsa04151 (PI3K-Akt signaling pathway), when they investigated the role of TCAB1 gene in the occurrence and development of head and neck carcinomas [48]. FBLIM1 was one of the genes corresponding to probes dysregulated in uninvolved oral samples and tumor samples of OSCC patients comparing to normal oral mucosa from non-cancerous patients [49]. Vincent-Chong et al. (2017) reported that SAMD12 gene was located in the cytoband 8q24.12, where copy number alternations identified by their study shared similarity with the TGCA of the oral cancer array CGH OSCC study [50]. Sun et al. (2014) reported that SESN3 gene was over-expressed after knock-down TCAB1 gene in head and neck carcinoma clinical specimens [48]. Baqordakis et al. found that FNDC1 proteins were up-regulated in OSCC [51]. The above observations indicated that the 258 genes corresponding to the 274 DV-only probes are relevant to OSCC and might provide new insights into the molecular mechanisms of OSCC. Further investigation and functional validation are warranted.

Our results showed that the number (1,639) of gene probes over-expressed in OSCC samples is not much different from (roughly 2 times) the number (736) of gene probes under-expressed in OSCC samples. However, the number (433) of gene probes over-variable in OSCC samples is much more than (roughly 34 times) the number (13) of gene probes under-variable in OSCC samples. This is consistent with what observed by Ho et al. [22].

The 5 papers mentioned in the Introduction section (i.e., Ho et al., 2008; Strbenac et al., 2016; Ran and Daye, 2017; Rahmatallah et al., 2017; and Sanati et al., 2018) did not compare their methods with the BF test. It would be interesting in evaluating the performances of the DV-detection methods used in those 5 papers with the BF test and other existing robust methods using systematic simulation studies and using real data analyses in future research.

There are some limitations in our study. The sample sizes of the two datasets that we used are relatively small. To guard against false positive discovery, we used one dataset as the discovery set and the other dataset as validation set and used robust equal-variance test. Based on the current knowledge, the 258 genes corresponding to the 274 DV-gene probes are not enriched in KEGG pathways. In future, we can perform functional validation in OSCC cases or cell lines to discover their functions and to validate their association with OSCC. Environmental factors, such as smoking, alcohol consumption, and HPV infection, commonly affect genetics in OSCC patients. In future, we will develop methods to incorporate these environmental factors in the investigations. HNSCC in the validation set might be different from OSCC. In future, more appropriate validation sets should be used.

## Acknowledge

This work is supported by China National Science and Technology Major Project with no.2017YFB1400803. The funding source had no direct role in study design, data collection, data analysis, in writing the report, or in the decision to submit the paper for publication

